# Association Study based on Topological Constraints of Protein-Protein Interaction Networks

**DOI:** 10.1101/2020.03.06.981175

**Authors:** Hao-Bo Guo, Hong Qin

**Affiliations:** Department of Computer Science and Engineering, The University of Tennessee at Chattanooga; SimCenter, The University of Tennessee at Chattanooga; Department of Biology, Geology and Environmental Science, The University of Tennessee at Chattanooga

## Abstract

The non-random interaction pattern of a protein-protein interaction network (PIN) is biologically informative but its potentials have not been fully utilized in omics studies. Here, we propose a network-permutation-based association study (NetPAS) method that gauges the observed interactions between two sets of genes based on the comparison between permutation null models and the empirical networks. This enables NetPAS to evaluate relationships, constrained by network topology, between gene sets related to different phenotypes. We demonstrated the utility of NetPAS in 50 well-curated gene sets and comparison of association studies using Z-scores, p-values or overrepresentations. Using NetPAS, a weighted human disease network was generated from the association scores of 19 gene sets from OMIM. We also applied NetPAS in gene sets derived from gene ontology and pathway annotations and showed that NetPAS uncovered functional terms missed by DAVID and other network-based enrichment tools. Overall, we show that NetPAS can take topological constraints of molecular networks into account and offer new perspectives than existing methods.

## Introduction

Interactomes, particularly, the protein-protein interaction networks (PINs) from model organisms and humans (Ito et al. 2001; Li et al. 2004; Rual et al. 2005; Stelzl et al. 2005; Hein et al. 2015; Huttlin et al. 2015; Huttlin et al. 2017; Li et al. 2017), have shifted our interests in molecular biology from the functions of individual genes or proteins to functional modules of PINs, including the modules associated with human diseases (Barabasi et al. 2011). PINs can be treated as graphs in which vertices (nodes) are proteins and edges (links) are protein-protein interactions. PINs possess characteristics observed in other real-world graphs, such as small-world (Watts and Strogatz 1998), scale-free (Barabasi and Albert 1999), and error-tolerance (Albert et al. 2000; Maslov and Sneppen 2002), suggesting that the topological patterns of PINs can offer biological insights.

Tools such as Gene Set Enrichment Analysis (GSEA) (Subramanian et al. 2005) and pathway analysis (Wang et al. 2007) have become a routine to extract shared characteristics of gene sets obtained from omics experiments. These analyses are often based on knowledge bases such as the gene ontology (GO) knowledge database (The Gene Ontology 2019), gene pathway databases such as KEGG (Kanehisa et al. 2019), and the interaction networks connecting genes or gene products (Barabasi and Oltvai 2004). Gene set (Huang et al. 2009a) (e.g., GSEA) and pathway (Khatri et al. 2012) analysis methods typically adopt statistical methods including Fisher’s, binomial, hypergeometric distribution, Chi-square, linear regression or logistic regression (Rivals et al. 2007; de Leeuw et al. 2016) to score the associations between the gene set and GO, pathways or other functional terms. Another work uses the cohesion coefficient to measure the association among pathways, annotations and gene sets (Yue et al. 2015; Yue et al. 2018). An implicit assumption of these analyses is that biological events in the cell are often conducted by groups of genes via direct, physical interactions, which are collectively called the interactome (Ghadie et al. 2018). In this regard, methods that are directly based on interactome, such as PIN, can take unique biological constraints into accounts and may offer more biologically relevant results than simple enrichment tests.

Multiple network-based approaches have been developed for functional predictions of gene sets. EnrichNet uses prioritization scores to expand the interested protein via random walks over the PIN (Glaab et al. 2012). WebGestalt unifies over-representation analysis (ORA), GSEA, and network topology-based analysis into a gene set analysis toolkit (Liao et al. 2019). Other tools including NET-GE (Di Lena et al. 2015) and pathfindR (Ulgen et al. 2018) aim to identify densely connected or functionally related subnetworks from the protein-protein interactions to strengthen the enrichment analysis. The PAGER database, on the other hand, has integrated gene-set, network, and pathway analysis (GNPA) data resources into a gene-signature electronic repository (Yue et al. 2015; Yue et al. 2018).

In addition to the above network-based enrichment tools and databases, another useful and important approach comes from comparisons with random networks (Newman et al. 2001; Maslov and Sneppen 2002; Maslov et al. 2004). However, random networks such as the Erdös-Rényi (ER) random networks (Erdös and Rényi 1959) do not have the power-law distribution of the node degrees observed in PINs and other natural networks (Barabasi and Albert 1999). As pointed out by Maslov and Sneppen (Maslov and Sneppen 2002), as well as Newman et al. (Newman et al. 2001), the meaningful permutations should have the node degrees preserved. Using the random (or null) networks with the preserved degree and/or additional lower-level topological parameters (Orsini et al. 2015), the higher-level attributes of the original network can be abstracted from statistical comparisons. In an outstanding work using the permutation null models, the hub nodes in both PIN and regulatory networks were observed to avoid the interactions with other hub nodes, and this observation was proposed to contribute to the stability of the networks (Maslov and Sneppen 2002). Inspired by this work we term the random (or null) networks with preserved node degrees as the MS02 null models (Qin et al. 2003).

In present work, we propose an approach to evaluate gene sets by comparing molecular networks with the MS02 null models, and we term this approach the network-permutation-based association study (NetPAS). We validated the usefulness with 50 well-curated gene sets and established consistency by using Z-score versus p-value and Jaccard-index. We also estimated an appropriate cutoff of Z-score to infer enriched or suppressed associations. A weighted human disease network was constructed using the NetPAS approach. Moreover, we showed that NetPAS can be applied in gene ontology and pathway enrichment analysis. We propose NetPAS is a useful tool to extract biological information stored in gene sets.

## Results

### Association Z-score of two gene sets

As illustrated in Figure 1, we can use the Z-scores to evaluate the over- or under-representation of interactions between two gene sets A and B—where Set A is a group of genes, e.g., genes associated with colorectal cancer, and Set B is another group of genes, e.g., genes associated with breast cancer. The gene IDs for both sets are obtained from OMIM (Amberger et al. 2019). The two sets share 3 genes. NetPAS first calculates the total number of edges (interactions) between set A and set B that appear in the original network—the human InWeb_IM PIN (Li et al. 2017) used in the present work (Fig 1b). Then by comparing with the numbers of edges from null network models (one example is in Fig 1d), a Z-score can be calculated as

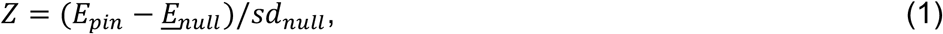

where *E*_*pin*_ is the number of interactions—i.e., number of edges— from the PIN, and *<u>E</u>*_*null*_ and *sd*_*null*_ are the mean and standard deviation of the numbers of interactions from the null models. We used 10k null models in the present work. For interactions between both sets, 51 are observed in the PIN (Fig 1c), compared to 25.4±4.8 observed in 10k null network models (one example is in Fig 1e), yielding an association Z-score of (51-25.4)/4.8 = 5.3. In Figure 1b, very few isolated interactions can be seen. In contrast, many isolated interactions can be seen in one example of a permuted null network model in Figure 1d. The contrast suggests that genes with single interaction tend to interact with genes with more connections in Figure 1c, which illustrates the importance of topological constraints in association tests. The details of the null model construction can be found in *Methods*.

**Figure 1.**
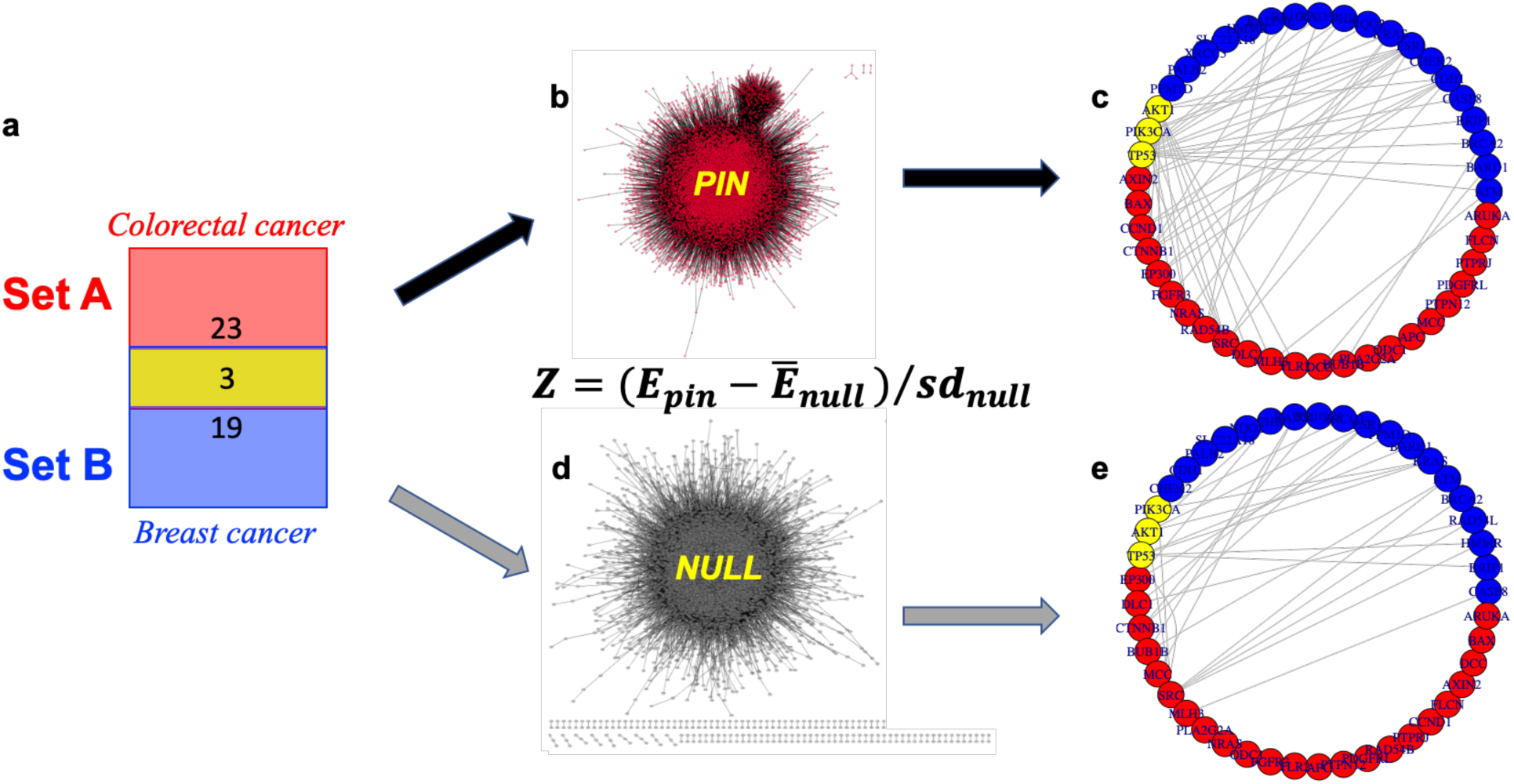
An example of calculating association Z-score of two gene sets. The two gene sets are selected from OMIM for colorectal cancer (MIM entry: 114500, set A) and breast cancer (MIM entry: 114480, set B). **a**) Venn diagram shows that Set A has 26 and Set B has 22 genes, respectively. There are three overlapping genes (*AKT, PIK3CA*, and *TP53*). **b**) The human PIN. **c**) 51 interactions between Set A and Set B observed from the PIN. **d**) An example of the null network model. **e**) 23 interactions between Set A and Set B observed from the null model shown in d. The interaction numbers from 10k null models are 25.4±4.8, leading to a Z-score of 5.3. Hence there is an enriched association between both cancers.

### Application in hallmark gene sets and the advantages of using Z-scores

We used 50 hallmark gene sets from the molecular signature database (MSigDB) (Liberzon et al. 2015). These hallmark sets can be considered “refined” benchmarks on top of >20k gene sets in MSigDB (version 7), which respectively represent well-defined biological processes with coherent expressions (Agarwal et al.). The names and details of these hallmark sets are listed in the Table S1 of the Supporting Information (SI). The gene names can also be found in Figure 2a, the boxplots of Z-score distributions of all hallmark sets. We calculated association Z-scores (eq. 1), one-tailed p-values and Jaccard-indices between all pairs of gene sets (including self-interactions). Figure 2b shows the heatmap of the association Z-scores calculated from all pairwise associations among the 50 hallmark gene sets using 10k MS02 null models compared with the original PPI. In this heatmap, positive Z-score (red) indicate over-representation, whereas negative Z-score (blue) indicates under-representation, respectively.

**Figure 2.**
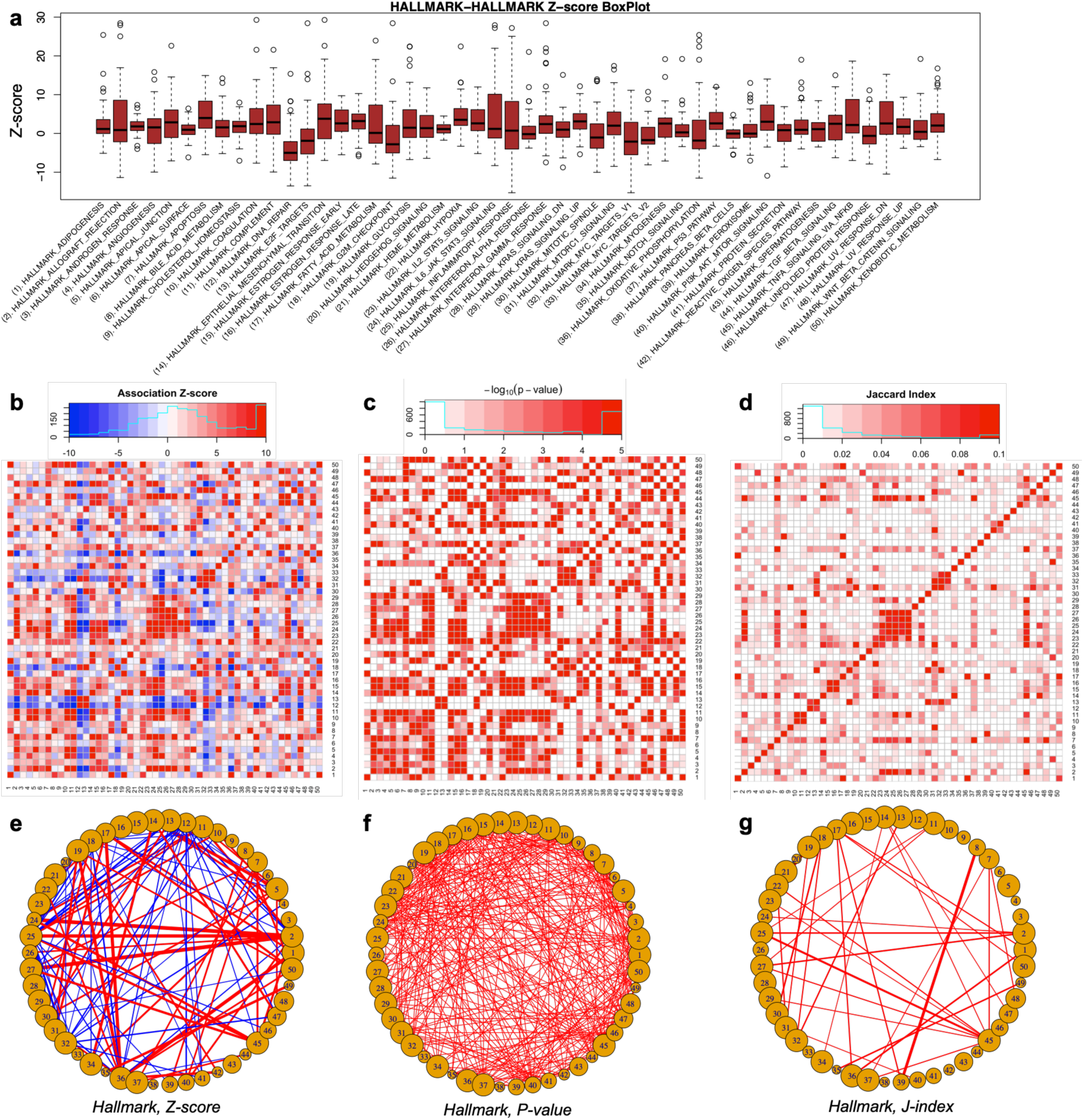
Association studies of 50 hallmark gene sets from MSigDB. (**a**) The boxplots of association Z-scores of the 50 gene sets with others—self-interactions are not considered in this plot. The names of the hallmark gene sets are listed along with their serial numbers. Heatmaps of (**b**) association Z-scores, (**c**) p-values, and (**d**) Jaccard-indices of the 50 gene sets as illustrated by their serial numbers. The names of the gene sets can be found in the Table S1 of the SI. Both enriched (red) and suppressed (blue) interactions can be revealed by Z-score. One-tail p-values (-log_10_ scale is used in the heatmap) are calculated for enriched interactions in the PIN compared to MS02 null models. When the observed interactions in the PIN are more than that in each of all 10k null models, we can only infer that P < 1e-4 instead of P = 0. We used P = 1e-5 (or −log_10_P = 5) for these situations. Networks of the hallmark gene sets have been generated for (e) Z-scores, the top 5% enriched (Z>11.8, red) and top5% suppressed (Z<-5.8, blue) interactions are shown in the network; (f) p-values, only those of P<1e-4 are have been shown, and (g) Jaccard-indices.

We directly calculated one-tailed p-values for associations from the PPI versus from 10k MS02 null models, based on a null hypothesis that the observed interactions are less than the null models. Heatmap of the p-values (log-scale) are plotted in Figure 2c and the p-value distribution is highly correlated with that of the Z-scores with a Pearson’s correlation coefficient (PCC) of 0.794 (p < 2.2e-16). The standard dissimilarity measure of Jaccard index J_AB_ is defined as J_AB_ = |A∩B|/|A∪B| for two gene sets A and B. J_AB_ is between 0 (0% overlap between A and B) and 1 (100% overlap between A and B). We also calculated the Jaccard-indices between the pairs of gene sets with a heatmap shown in Figure 2d. A general agreement was also observed between association Z-scores and Jaccard indices with PCC = 0.48 (p < 2.2e-16).

For comparison, we constructed networks to illustrate association patterns among the 50 hallmark sets using the association Z-scores, p-values, and Jaccard-indices, respectively. All networks use the gene sets as nodes and association scores as edge weights. Figures 2e, 2f and 2g show parts of all three networks, respectively. In the Z-score network (Figure 2e) top 5% over-represented (red) interactions have Z larger than 11.8, and the top 5% under-represented (blue) interactions have Z smaller than −5.8, respectively. The p-value network (Figure 2f) shows 326 associations (for all 1225 pairs of gene sets excluding the self-interactions) with p-value < 1e-4 (i.e., more interactions observed in the PPI than all null models), and in this network a uniform edge-weight is applied for these interactions. For comparisons, the Z-score network in Figure 2e has 76 positive and 61 negative interactions, the p-value network (Figure 2f) has 326 interactions, whereas the Jaccard network (Figure 2g) has only 40 interactions, respectively. Note that all networks would possess more interactions by using looser cutoffs. For instance, a criterion of |Z|>2 for the Z-score network would lead to 509 positive and 254 negative interactions; a cutoff of p-value <1e-3 results in 427 interactions; and a cutoff of J > 0 for the Jaccard network—similar to a previous human disease network (Goh et al. 2007) in which two diseases are connected if they share at least one gene—would lead to 871 interactions, respectively.

Estimations of p-values are limited by the number of null models used. For 10k null models applied in present work, we cannot validate a p-value smaller than 1e-4, which is roughly equivalent to Z=3.72 for a one-tail test under normal distributions. Therefore, based on limited number of null models, it is difficult to rank the interaction strengths that have low p-values to a graph, as such, in Figure 2f p-value <1e-4 interactions are visualized with a uniform weight. However, the Z-scores (Figure 2e) spread a considerably wide range using a limited number of null models. In addition, we show that in a Z-score heatmap (Figure 2b) or network (Figure 2e), both enriched (red) and suppressed (blue) associations can be plotted. However, for the one-tail p-value analysis, only one of both associations can be addressed at a time, based on the null hypothesis used—such as enriched associations in Figure 2c and 2f—despite both enriched and suppressed associations can be analyzed separately. Similar to using p-values, the Jaccard indices also cannot describe the under-representation information on how gene sets ‘avoid’ interacting with each other and is only informative on enrichment. Using Z-scores we can identify both enriched and suppressed interactions with relatively small number of null network models: for example, the standard deviation of the Z-score differences between using 10k and using 1k null models is 0.146, which is considerably smaller than potential cutoffs, e.g., 11.8 for the top 5% or −5.8 for the bottom 5% Z-scores, or less than 10% of the cutoff of |Z|=2, see below in the discussion of random models.

Interestingly, for all 326 interactions with p-value <1e-4, their Z-scores are 9.59±6.36 with Z_min_ = 3.78, which are equivalent to p-value <1e-4 under a normal distribution one-tail test. Moreover, 11 of these 326 interactions have Jaccard-index of 0: although these 11 pairs show positive interactions (p-value <1e-4 and Z=6.11±1.97 with Z_min_=4.24), no shared genes between each pair could be found. One example is for gene sets 6 and 24 (full names in Table S1 of SI) that have J=0, p-value<1e-4, and Z=10.3, which reflects a significant over-represented interaction number (344) in the PIN compared to null models (204.5±13.5), as shown in Figure S1 of the SI.

A negative Z-score calculated by NetPAS reflects under-represented interactions between two gene sets. In the box plot of Z-scores between gene sets (self-interactions are excluded, Figure 2a), some hallmark sets appear to have a negative mean Z-score and appear to have ‘avoided’ interactions to most of the other hallmark sets. For example, the hallmark set 12 (full name in Table S1 of SI) has a mean Z-score of −3.6 for interactions with all other hallmark sets. Figure S1 in the SI shows interactions between set 12 and set 25 observed from the PIN and a representative null model.

### Recommended cutoffs for application in practice based on background Z-scores

To find out recommended Z-score cutoffs for application of NetPAS in practice, we constructed random gene sets with comparable sizes to MSigDB in Figure 3a (Figure S2 of SI). The association Z-scores among these random sets are narrowly centered around zero (color bar in Figure 3a). In contrast, association Z-scores of the 50 hallmark sets have a long-tailed distribution with a skewed-peak at the positive upbound (color bar in Figure 2a). The association Z-scores between the random gene sets (Figure 3a) are much less and looser than the hallmark gene sets. Moreover, as randomly constructed networks reflect the genetic background, distributions of the Z-scores among these random gene sets can be used to validate the cutoffs for quantifying associations of gene sets. For all 15k association Z-scores between random gene sets, 451 have Z > 2 and 160 have Z < −2, corresponding to p-value = 0.030 and p-value = 0.011, respectively (Figure 3b). Self-associations are excluded in Figure 3b although they do not show noticeable differences to non-self-associations for the random sets. For the random gene sets, using |Z| > 2 as a cutoff we observed a limited number of enriched (red, 30) and suppressed (blue,) associations (Figure 3c), which are much less than the hallmark gene sets. Similar trends are found in random networks of different sizes (Figure S2 of SI).

**Figure 3.**
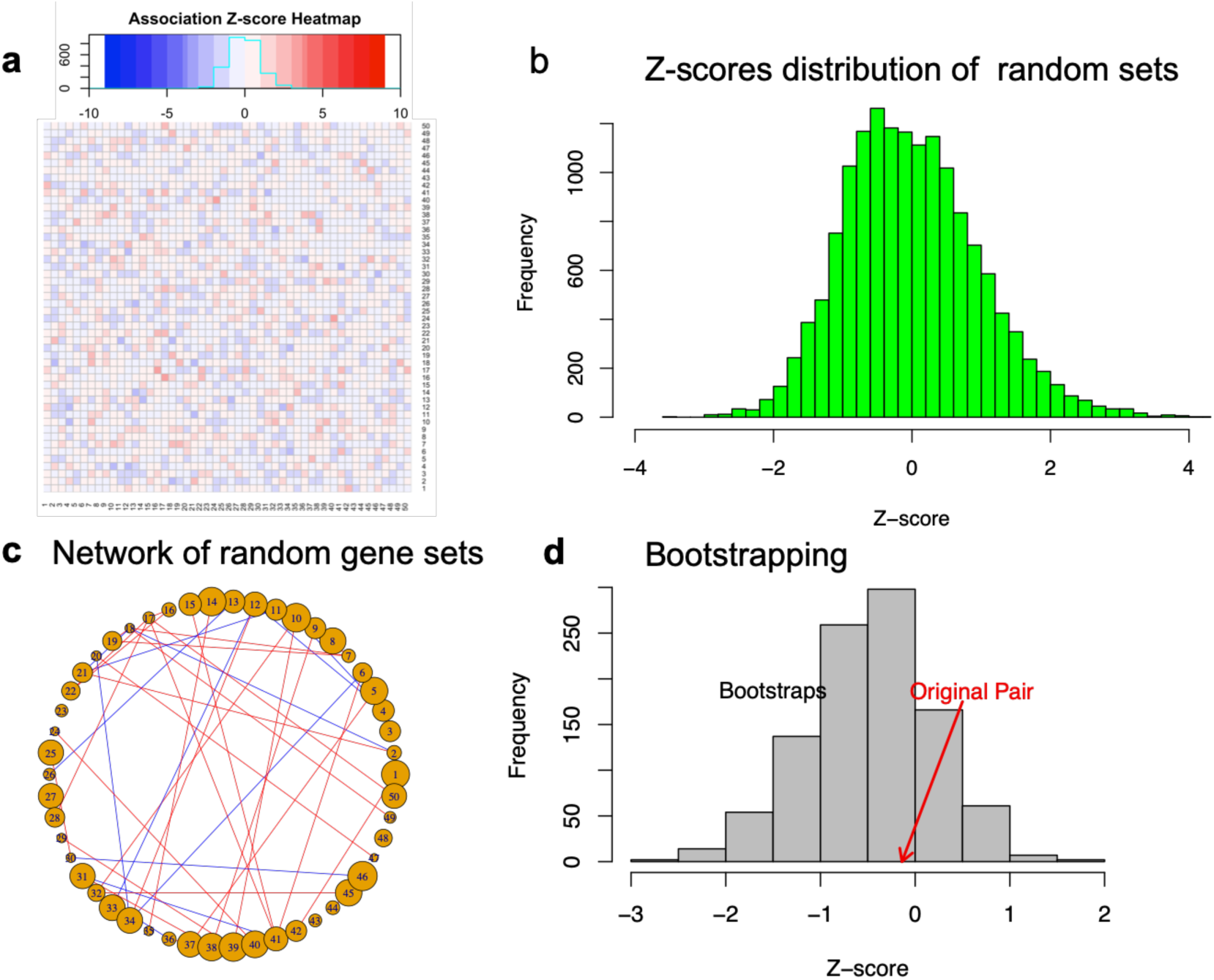
Association Z-scores of random gene sets and statistical validation of cutoffs. (**a**) Heat map of association Z-scores among 50 randomly constructed gene sets with gene numbers in the range of [15,200]. (**b**) Histograms of association Z-scores calculated from five sets of random networks (see Methods and Figure S2 in the SI). Among all 15k Z-scores between random gene sets, 451 have Z>2 (P = 0.030) and 160 have Z<-2 (P = 0.011), respectively, indicating that using |Z|>2 as the cutoff would be appropriate. (**c**) Network of the random gene sets weighted by the association Z-scores. A cutoff of |Z|>2 is used for both enhanced (red) and suppressed (blue) interactions. (d) We then bootstrapped the affiliations of all genes from the two randomly constructed gene sets, each contains 100 genes. The association Z-score of the original pair is −0.14 (red arrow) whereas the Z-score of the 1000 bootstraps are −0.47 ± 0.66.

The association Z-score between two gene sets—say, set A and B—reflects how likely the genes in set A favor (Z>0) or avoid (Z<0) the interactions with those in set B, and vice versa. Note that although for normal distributions |Z| > 2 is roughly equivalent to p < 0.023 from a one-tailed t-test; however, the normal distributions can hardly be applied to power-law networks including PPIs, and different cutoff in the Z-scores may lead to different interpretations. Nevertheless, we observed that compared to randomly constructed gene sets a cutoff of |Z| > 2 is appropriate (Figure 3b).

To further understand how to interpret association Z-scores, we selected two randomly constructed gene sets, each comprises 100 genes, that have no apparent association with Z = −0.14. In this example the number of the bootstrap combinations is 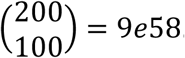. We did not sample all bootstraps. Instead, using 1000 bootstraps, the Z-scores are distributed in −0.47±0.66, as shown in Figure 3d. The ratio of |Z| > 2 is 0.016 for all bootstraps (i.e., p=0.016 for a two-tail test). Therefore, we suggest that in practice, a cutoff of |Z| > 2 is appropriate, in line with the discussions of the random constructed gene sets as shown in Figure 3b.

The above analysis indicates that the background association of gene sets has relatively small Z-scores and it is an appropriate practice to use a cutoff such as |Z| > 2 to infer an association between two gene sets.

### Constructing a weighted human disease network

A previous work (Goh et al. 2007) analyzed more than a thousand of human disorders with associated genes maintained by OMIM (Amberger et al. 2019). This work produced the “human disease network” (HDN), assuming that two disorders are connected if they share at least one gene, i.e., the Jaccard-index > 0. It was shown that the genes associated with the same disorder have a 10-fold increase of likelihood to interact with each other than those that are not associated (Goh et al. 2007).

Here, we use NetPAS to estimate the association Z-scores of 19 descriptive entries from OMIM. These entries are associated with different disorders and contain at least 5 associated genes for each entry. These entries include 13 cancers, 3 mental disorders and 3 other disorders (Table S2 of the SI). Although there is no association between certain diseases, such as Alzheimer’s and colorectal cancer shown in Figure 3D, the associations between some diseases are significant. The Z-score heatmap and the resulting weighted human disease network (wHDN) are shown in Figure 4. This wHDN has several isolated nodes, including esophageal cancer, renal cell carcinoma (RCC, a type of kidney cancer), pheochromocytoma (Pheoch, rare cancer related to the adrenal gland), Alzheimer’s and Parkinson’s diseases. Each isolated node contains 5-8 genes. However, some nodes with similar sizes are strongly associated to other diseases, such as ovarian cancer (6 genes), non-Hodgkin Lymphoma (NHL, 5 genes) and meningioma (6 genes). Therefore, the strength of associations between gene sets (disorders in this example) is not determined by the number of genes. Instead, the direct interactions between genes associated with the gene sets (disorders), and with comparisons to those observed in null network models, have contributed to determining the association strength of two gene sets.

**Figure 4.**
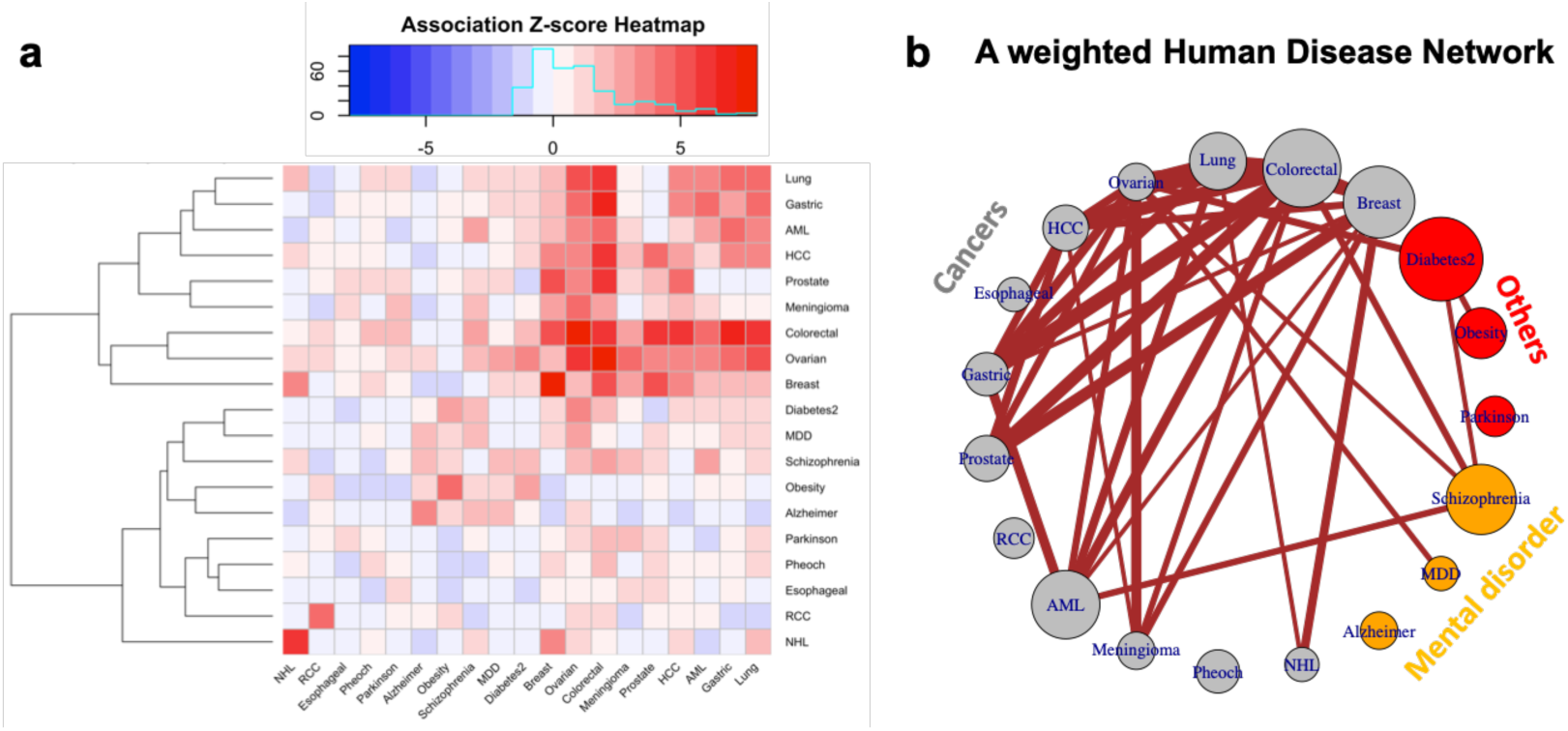
A weighted human disease network (HDN) generated by NetPAS. (**a**) The association Z-score heatmap of 19 diseases, which include 13 cancers (grey), 3 mental disorders (orange) and three other diseases (red). (**b**) A weighted human disorder network constructed from the Z-score matrix indicates that most cancers highly interacted with each other. Some mental disorders including Schizophrenia and Major depression disorder (MDD) are associated with certain cancers, and Type II Diabetes (Diabetes2) is associated with ovarian cancer. Note that in this wHDN all interactions are positive, and no negative or suppressed interactions among these diseases (i.e., Z<-2) have been observed.

The wHDN shown in Figure 4b indicates that 10 out of 13 cancers (except three isolated cancers mentioned above) have strong associations with each other. The mental disorders Schizophrenia and Major Depression Disorder (MDD) are highly associated with certain cancers, whereas Alzheimer’s is not. The Type-II Diabetes is also associated with cancer as well as Schizophrenia. Obesity is not directly associated with cancer but is associated with Type-II Diabetes. This result may be useful to the understanding of disease-disease relationships. In summary, NetPAS is useful to evaluate the associations among diseases based on physical interactions, instead of overrepresentations of genes.

### GO and pathway enrichment analyses using NetPAS

A GO term or a pathway functional term can be regarded as a gene set affiliated to this term. Because NetPAS can be used to estimate the association strength between any two gene sets, it is straightforward to extend the NetPAS approach to the GO and pathway enrichment analysis. For a given target gene set, its association Z-scores with all gene sets related to the GO/pathway functional terms can be separately calculated and ranked, from which the enriched or suppressed functional terms can be inferred. To demonstrate this utility, we performed the GO (The Gene Ontology 2019) term and KEGG (Kanehisa et al. 2019) pathway enrichment analysis of the 50 hallmark gene sets (see above), and compared the results with those obtained by a traditional enrichment method DAVID (Huang et al. 2009b). In this analysis, the association Z-scores are calculated between the target gene set and the 18,033 gene sets derived from 17,715 GO terms and 318 KEGG pathways (see Methods). All GO terms and KEGG pathways are then ranked to infer both enriched and suppressed functional terms.

The top 10 enriched terms by both NetPAS and DAVID for one example, *HALLMARK_OXIDATIVE_PHOSPHORYLATION*, were shown in Figure 5a. Consistency between the two methods can be seen: 9 out of 10 BP terms, 10 out of 10 CC terms, 8 out of MF terms, and 8 out of 10 KEGG terms predicted by NetPAS are also predicted by DAVID. However, some functional terms detected by NetPAS are missed by DAVID and other enrichment tools, such as the BP term GO:0015990 (“electron transport coupled proton transport”). In this example, the target hallmark gene set has 94 interactions with genes that carry the term GO:0015990, observed in the PIN. In contrast, there are only 4.7±2.1 interactions from the 10k null models, leading to a large Z-score of 42.9 for this GO term. For all 50 hallmark sets and the top-10 enriched GO terms by NetPAS, 73.4% BP, 70.2% CC and 55.0% MF terms were verified by DAVID. For all functional terms suggested by NetPAS but missed by DAVID, the enrichment signals come from the fact that more interactions between the target set and the function annotation term have been observed in the PIN than random null models. Figure 5b shows the subnetworks for interactions between the hallmark set exemplified in Figure 5a and the gene sets affiliated with the top-10 BP terms by NetPAS.

**Figure 5.**
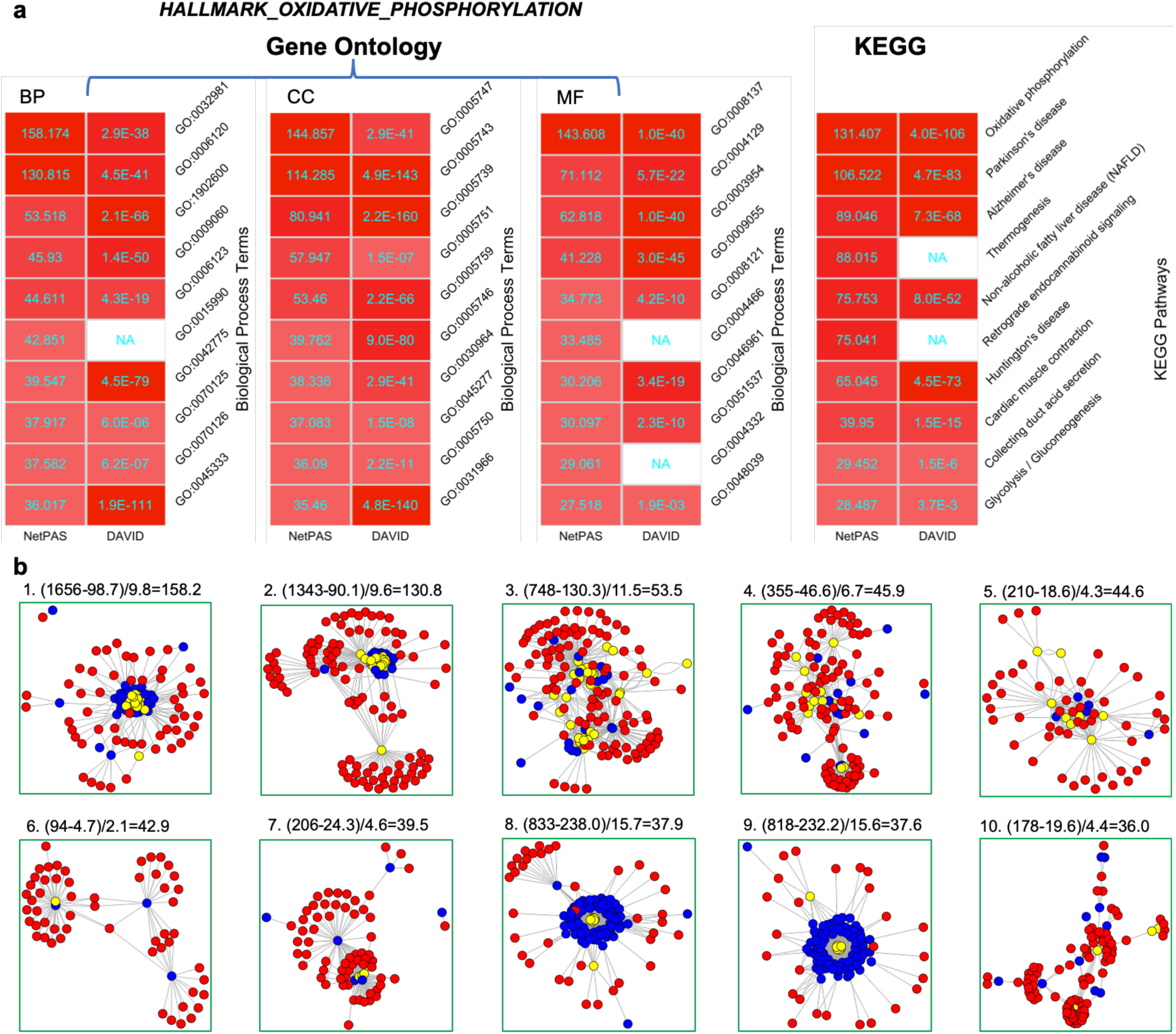
An example of GO and pathway enrichment performed by NetPAS. using the hallmark set *HALLMARK_OXIDATIVE_PHOSPHORYLATION*. (**a**) The Top ten enriched gene ontology terms, including biological process (BP), cellular component (CC) and molecular function (MF) (left), and enriched KEGG pathway terms (right) using NetPAS (left) and DAVID (right). The magnitude of enrichments is scaled by the colors from white (Z=0) to red (Z=Z_max_). The p-values estimated by DAVID were converted to Z-scores using a two-tailed normal distribution for coloring. (**b**) Interaction sub-network between the target gene (red nodes) and the genes affiliated with the top 10 biology process (BP) GO terms (blue nodes); the genes that are both affiliated with the functional term and belong to the target gene set are shown in yellow nodes. Formulas used for calculating the Z-scores for each BP term are written on top of each subnetwork.

As a network-permutation-based approach, NetPAS is sensitive to the subnetwork within the gene sets, including its global cluster coefficient, maximal cluster size, and maximal clique degree, summarized in Table S3 of SI. S6 Figure exemplified other gene sets from MSigDB with comparisons to DAVID and the network-topology-based method WebGestalt. We suggest that NetPAS can serve as a useful complementary tool to traditional as well as other network-based enrichment methods.

## Discussion

No gene or protein functions alone. The cellular functions can be regarded as being conducted by functional modules or communities (Hartwell et al. 1999) of genes/proteins in the interactome (Ghadie et al. 2018). The concept of disease module (Barabasi et al. 2011; Menche et al. 2015) has also been proposed based on the fact that the genes associated with the same disease are more likely to interact with each other (the “local” and “disease module” hypotheses in (Barabasi et al. 2011)). This principle can also be applied to other curated gene sets such as those in MSigDB (Liberzon et al. 2011; Liberzon et al. 2015). Indeed, for the 50 hallmark sets (Figure 2), the mean association Z-score excluding self-interactions (Figure 2d) is 2.0. However, the self-interactions for all gene sets have a significantly higher mean Z-score of 17.8. This trend holds for the 19 diseases shown in Figure 5. For the random sets shown in Figure 3, however, the mean Z-score is −0.04 and the mean self-association Z-score is −0.20. Therefore, our results indicate over-presented interactions for genes in the curated data sets, such as those from MSigDB or related to diseases, in contradict with random chances.

A biological network such as PIN is scale-free with the degrees of all nodes following the power law. Because in the null models of present work all node degrees have been preserved, they have the same power-law distribution as those in the original PIN. In biological networks, low-degree nodes tend to connect to high-degree nodes, or hubs (Maslov and Sneppen 2002). For example, there are 1004 nodes in the PIN with the degree k = 1, i.e., each of them only has a single interaction. In the PIN, only two interacting pairs (*CLEC2A*:*KLRF2* and *REC114*:*MEI4*) are formed by such nodes, constituting two isolated interacting pairs. However, for 10k null models, there are 113.3±47.6 isolated pairs with a minimum value of 13. Figure 6 in the Methods shows the histograms of the number of isolated pairs in all 10k null models, and an example is shown in Figure 1C.

**Figure 6.**
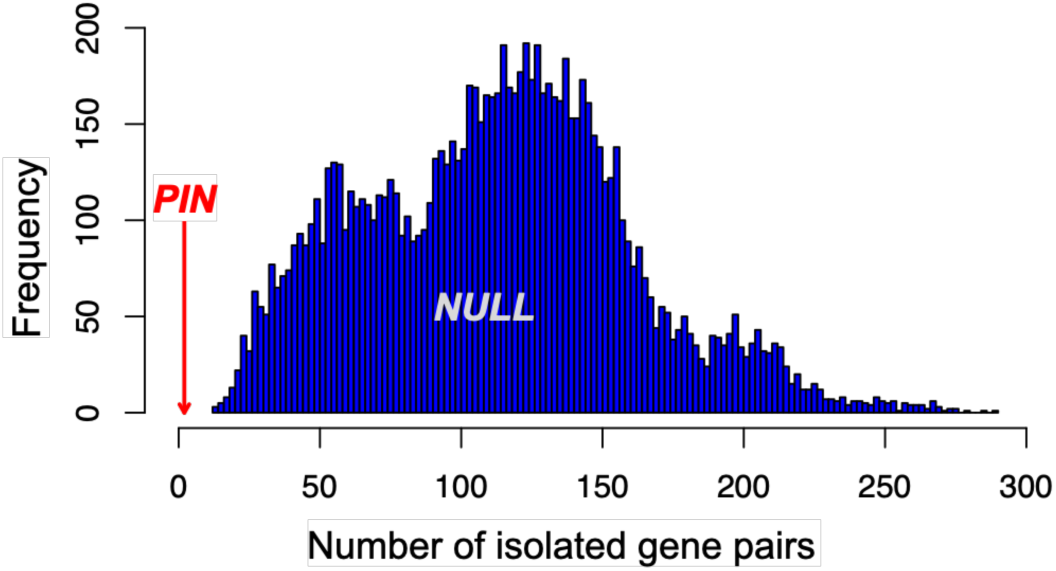
Topological differences between the original PIN and null models. exemplified by the number of isolated pairs (interactions between k=1 nodes) in the PIN with observation (red arrow) of 2. However, the number of isolated pairs in 10k null models is distributed from 13 to 289 with a median value of 115 (blue histograms). For comparison also see Figures 1b and 1c. The number of isolated pairs in the null models is significantly larger than that in the PIN (p<1e-4, for 10k null models).

Several limitations of NetPAS need to be emphasized, however. The first limitation is the incompleteness of the resources including the interactomes, the coverage of genes in different gene sets, and gene annotations in GO or pathway knowledge databases. In addition, protein-protein interactions are dynamic (Han et al. 2004), and they may vary significantly among different tissues or cell types (Ellis et al. 2012; Yao et al. 2018). These limitations may be addressed in future studies by the integration of tissue-specific or cell-type-specific interactomes to further our understanding of the biological significances of different gene sets.

In summary, we show that NetPAS can quantify the association between two different gene sets by taking network constraints into account. We demonstrate the utility of using Z-scores in NetPAS compared to using p-values or Jaccard-scores. NetPAS is useful in classifications of gene sets, including those associated with different diseases. We also show that NetPAS can be applied in GO and pathway enrichment analysis, in which every single GO or pathway functional term is regarded as an affiliated gene set. The NetPAS approach can be applied to extrapolate the biological association between different gene sets such as potential relationships between various gene sets behind different phenotypes and diseases. NetPAS can also be applied in other types of networks to estimate the association strengths between network subsets.

## Methods

### MS02 null permutation of the PPI network

The permutation-based network null model is based on a work of Maslov and Sneppen in 2002 (Maslov and Sneppen 2002) (hence named MS02 null model in present work). The human PIN used in the present work contains 592,685 edges spreading on 16,641 nodes. This PIN is considered as simple graphs, i.e., it is undirected and does not contain self-interactions (self-loops) or multi-interactions.

A network is regarded as a graph *G* = (*V, E*) with order of |*G*| = *N*, the vertices (or nodes) are *V* = {*v*_1_, *v*_2_, … *v*_N_}. An MS02 null model, *G*^null^ = (*V*^null^, *E*^null^), has

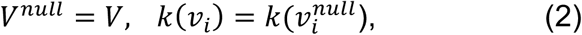

where *k*(*v*_*i*_) is the degree or edge numbers—also known as connectivity—associated with *v_i_*, i.e., it uses the same vertex set and degrees for all vertices are preserved as the original network.

It worth noting that all MS02 null network models obey the power-law because the node degrees are the same as the observed PIN. However, there are significant topological differences between the null models and the PIN. For example, in the PIN there are only two isolated interactions (*CLEC2A*:*KLRF2* and *REC114*:*MEI4*) connected by the nodes with degree k = 1. However, for 10,000 null models, there are 113.3±47.6 isolated pairs with a minimum value of 13 connected by the k = 1 nodes (Figure 6). The abundance of isolated interactions in MS02 null models indicates that the power-law distribution of node degrees does not originate that the low-degree vertices tend to connect to the hub vertices, as suggested previously (Barabasi and Albert 1999). Instead, the low-degree vertices tend to interact with the high-degree vertices may be a unique feature of the natural networks such as the PINs, compared to MS02 null network models.

### Protein-protein interaction network (PIN)

Human protein symbols of both the PIN and protein sets (see below) adopt the HUGO gene nomenclature (Braschi et al. 2019). The InWeb_IM network (v 2016_09_12) (Li et al. 2017) was taken for the human PPI network. This network does not allow self-loops (as the MS02 models) and comprises 592,685 edges (downloaded in August 2018). InWeb_IM is one of the largest protein physical interaction networks; e.g., 1.8x of a recent release of the BioGIRD (Oughtred et al. 2019) human PIN (v3.5.168, 326,529 physical-interaction edges). Importantly, this network showed excellent performance in representing the gene-gene relationships across hundreds of human pathways (Li et al. 2017; Li et al. 2018), as well as in assisting the discovery of genes associated with diseases such as cancers (Horn et al. 2018), which is particularly suitable to the goal of present work.

### Gene Ontology (GO) Annotations and Pathways

All GO annotations were downloaded from the Gene Ontology Consortium website (www.geneontology.org) (Ashburner et al. 2000; The Gene Ontology 2019) updated February 2019. The GO terms are grouped in three basic ontologies: biological process (BP), molecular function (MF) and cellular component (CC). The GO annotations for human genome include 11,883 BP terms for 17,697 genes, 4,128 MF terms for 17,498 genes, and 1,704 CC terms for 18,697 genes, respectively. The KEGG (Kyoto Encyclopedia of Genes and Genomes) pathways have been obtained from the ConcensusPathDB (Herwig et al. 2016). Altogether 318 KEGG pathways for 7,344 genes have been used.

For all 50 hallmark gene sets from MSigDB, we used the conventional enrichment tool DAVID (Huang et al. 2009b) for the GO and pathway enrichment analysis. We also used the network-topology-based tool WebGestalt (Liao et al. 2019) for the predictions, see Figure S3 in the SI.

### Gene Sets

Fifty sets of human proteins from the MSigDB hallmark gene set (category H) collection (Liberzon et al. 2011; Liberzon et al. 2015) were used to evaluate the interactions between gene sets. 19 gene sets associated with different human disorders were obtained from OMIM (Amberger et al. 2019), including 13 cancers, 3 mental disorders, and 3 other diseases, as listed in the Table S2 of the SI.

We also randomly constructed six gene-set groups, each of which comprises 50 gene sets. Gene sets in groups I, II, III, and IV have the same number of 15, 50, 100 and 200 genes for each set, respectively. For each gene set in group V, the gene number is randomly set in the range of [15,200], and for each gene set in group VI, the gene number is randomly set in the range of [1,100]. Each group has 50 self-associations and 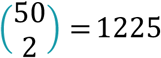 unique non-self-associations. The distribution of Z-scores of these associations are shown in Figure 3.

All calculations are performed using R. The resources and simulation codes are deposited in the GitHub repository at https://github.com/QinLab/NetPAS.

## Acknowledgments

This work is supported by NSF Career award #1453078 (transferred to #1720215), BD Spoke #1761839, and internal funding of the University of Tennessee at Chattanooga. All simulations are performed using the SimCenter computing resources of the University of Tennessee at Chattanooga.

## Author contributions

H.B.Guo and H.Qin conceived the study, designed the workflow, performed data analysis and wrote the manuscript.

## Additional information

**Supplementary information** is available for this paper.

